# Multidimensional Analysis of Bronchoalveolar Lavage Cytokines and Mast Cell Proteases Reveals Interferon-γ as a Key Biomarker in Equine Asthma Syndrome

**DOI:** 10.1101/2020.02.24.956573

**Authors:** Jane S. Woodrow, Melissa Hines, Carla Sommardahl, Bente Flatland, Kaori U. Davis, Yancy Lo, Zhiping Wang, Mary Katherine Sheats, Elizabeth M. Lennon

**Author notes:** Address correspondence to (EML).

## Abstract

Naturally-occurring equine asthma is an inflammatory lung disease characterized by chronic, partially reversible airway obstruction, pulmonary remodeling and lower airway inflammation. The cytokine profiles that distinguish asthma groups or subtypes in horses have not been systematically classified, and mast cell phenotypes, which, in human asthma, correlate with asthma type, lung function, and response to therapy, have not been well-described in horses. The purpose of this study was to: (1) compare mast cell protease mRNA expression between healthy and asthmatic horses, (2) analyze the cytokine profile present in BALF of currently defined equine asthma groups, and (3) use these data to evaluate potential biomarkers of defined asthma groups. Mast cell protease gene expression and select cytokine gene expression in cells isolated from BALF, and BALF multiplex cytokine assays were performed. Multidimensional analysis demonstrated that IFNγ differentiates severe from moderate asthma, and that TNFα and CXCL8 are key biomarkers of equine asthma subtype. Expression of chymase mRNA, a mast cell-specific protease, was significantly decreased in horses with mastocytic asthma. These results will help further define EAS immunopathology, which could improve understanding and definitions of asthma groups, while also potentially identify novel therapeutic strategies.

## Introduction

Equine asthma is a chronic inflammatory lung disease characterized by enhanced bronchial reactivity, and chronic, partially reversible airflow obstruction, pulmonary remodeling, and lower airway inflammation [1]. The most common environmental triggers for asthma development in susceptible horses include pollens, dust, mold, and bacterial components. Equine asthma prevalence in the Northern hemisphere is about 14% and the disease can be frustrating to manage, while also being career or life ending for the horse [2, 3]. Recently it has also been shown that prevalence can change with season [4]. The diagnosis is based on history, physical examination, bronchoalveolar lavage fluid (BALF) cytology, and lung function tests. Lung function tests are performed less frequently due to cost, required skill for equipment use and data analysis, and availability of the apparatus. History and physical exam findings supporting asthma include clinical signs of lower airway inflammation, such as cough, increased respiratory rate and effort, poor performance/exercise intolerance, and serous, mucoid, or mucopurulent nasal discharge at rest and/or at exercise. A 2016 consensus statement has defined two categories of equine asthma syndrome (EAS), currently designated as mild to moderate asthma (mEAS) and severe asthma (sEAS) [1]. For horses with sEAS, auscultation of the lungs at rest or during rebreathing exam can reveal wheezes, crackles, cough, and/or tracheal rattle. Abnormalities upon auscultation may not be apparent, but recovery from a rebreathing exam may be prolonged in mEAS or sEAS horses. In contrast to horses with sEAS, horses with mEAS do not have clinical signs at rest [1]. BALF cytological findings supporting asthma include an increased percentage of neutrophils (most pronounced in severe asthma), mast cells, eosinophils, or some combination of increase in these cells.

In both horses and humans, asthma immune response phenotypes have been broadly divided into allergic asthma, with a Th2-high immune response, and non-allergic asthma, with a non-Th2 immune response. However, the two phenotypes are not always clearly defined as such and can include horses from both mEAS and sEAS group [5]. A Th1 immune response has been shown to be associated with a generalized increase in BALF cell numbers in lower airway inflammation, although cytokine profiles, either at the protein or mRNA level, may support more of a Th1, Th2, Th17, or mixed immune response, depending on the study [6–17]. Additionally, the extent of airway remodeling and collagen deposition has been linked to the balance of transforming growth factor (Tgf)-β1 and bone morphogenetic protein (BMP)-7 in human and mouse asthma [18–20]. One report in horses notes no significant differences in Tgfβ1, but has not been followed up, nor has BMP7 been investigated [21]. Due to inconsistencies in reported cytokine profiles and focusing on a single category of EAS, this has limited our ability to fully describe the pathogenesis of EAS and its categories, which ultimately limits our diagnostic, treatment, and management abilities.

Asthma in people is additionally described by the type of inflammatory cells present in the airways [22–24]. Recently, the presence of mast cells in human asthmatics’ BALF and induced sputum samples has been shown to correlate with asthma type and response to therapy [25–27]. In humans, two main phenotypes of mast cells are recognized based on the proteases they contain; tryptase-only containing mast cells (MC_T_), and tryptase, chymase, and carboxypeptidase A3 containing mast cells (MC_TC_) [28, 29]. Healthy human lungs predominantly contain mast cells of the mucosal-type (MC_T_), and asthma is associated with an increase in MC_T_ and/or the connective tissue-type (MC_TC_) depending of asthma type [25, 30]. Cytological identification of mast cells in induced human sputum is difficult due to low cell prevalence; therefore, protease mRNA expression is used. Using this technique, a third mast cell type with increased tryptase and carboxypeptidase A3, but decreased chymase expression was identified in human asthmatics that correlated with eosinophilic asthma and increased responsiveness to corticosteroids [26, 27]. Mast cell lung phenotypes identified in humans have not been thoroughly evaluated in horses, but warrant further investigation as they are limited to tissue samples [31, 32].

We hypothesized that healthy and asthmatic horses would have significantly different BALF mast cell protease mRNA expression and cytokine/chemokine levels that would correlate with BALF cytology results and health status. The objectives of this study were to: (1) compare mast cell protease mRNA expression between healthy and asthmatic horses, (2) analyze the cytokine profile present in BALF of currently defined equine asthma groups, and (3) use these data to evaluate potential biomarkers of defined asthma groups.

## Results

### Main study population – University of Tennessee College of Veterinary Medicine (UT-CVM)

A total of 54 horses were sampled for evaluation. Twenty horses were excluded because they did not meet our inclusion criteria (see **Table 1 and Supplementary Table S3**). Final groups were comprised of 11 healthy and 23 asthmatic horses (9 mild/moderate and 14 severe) (see **Supplementary Table S1 and S2 online)**. Median age was not significantly different between groups. Various breeds were represented in each group.

**Table 1.** Diagnostic Criteria for Experimental Groups.

| **Experimental Group** | **History & Physical exam** | **CBC/fibrinogen** | **BALF cytology** |
| --- | --- | --- | --- |
| **Healthy**<br>*Main population*<br>*UT-CVM* | Normal | Normal | ≤6% neutrophils,<br>≤1% eosinophils,<br>≤2% metachromatic<br>cells, and<br>≤50% lymphocytes |
| **Asthma**<br>*Main population*<br>*UT-CVM* | History, clinical signs, and<br>physical exam supportive<br>of lower airway<br>inflammation; no changes<br>to or initiation of<br>corticosteroids (systemic<br>or intranasal) within 2<br>weeks of enrollment | Not consistent<br>with systemic<br>infection | ≥10% neutrophils,<br>≥5% mast cells,<br>and/or ≥5%<br>eosinophils<br>*severe asthma,<br>>25% neutrophils |
| **Mastocytic Asthma**<br>*Sub-population*<br>*NCSU-CVM* | History, clinical signs, and<br>physical exam supportive<br>of lower airway<br>inflammation; no<br>corticosteroids<br>administered within 2<br>weeks of BAL | Not performed | ≥2% mast cells<br>≤6% neutrophils<br>≤1% eosinophils |

### Sub-population – North Carolina State University College of Veterinary Medicine (NCSU-CVM)

Samples from 8 horses with mastocytic asthma (maEAS) sampled at NCSU were included to fully evaluate mast cell protease expression due to the lack of mast cells detected in UT-CVM asthma cohort (see **Table 1, Supplementary Table S1, and S2 online**). Although the ages of some of the horse were unknown, the median age of the NCSU-CVM group was not significantly different from the UT-CVM groups.

### BALF Cell Count and Cytologic Analysis (UT & NCSU-CVM)

Results of BALF cell count and cytological analysis are described in **Table 2** and **Supplementary Table S2**. The median total nucleated cell count (TNCC) of each group was not statistically different, although 5 outliers in the sEAS group had TNCC > 1,000 cell/μL. As expected, based on standard diagnostic criteria, horses with severe asthma had significantly higher percentages of neutrophils compared to healthy horses and those with maEAS (p < 0.0001 & p = 0.0002, respectively). Horses with sEAS had lower percentages of lymphocytes in their BALF when compared to mEAS or those with maEAS (p < 0.0001 & p = 0.0009, respectively). The composition of immune cells in healthy horses was predominated by macrophages compared to sEAS (p < 0.0001). Three horses with mEAS had BALF eosinophilia with ≥ 5% eosinophils. No eosinophils were detected in BALF from any healthy horses or horses with sEAS. Mast cells were only detected in maEAS horses with a median mast cell percentage of 5.0 [interquartile range (IQR): 4.18 – 5.75%].

**Table 2.** Bronchoalveolar lavage fluid characteristics for the four defined groups: TNCC, Neutrophil %, Lymphocyte %, Macrophage %, Eosinophil %, and Mast cell %. Data are expressed as median [IQR]. sEAS, n=14; mEAS, n=9; Ctr, n=11; maEAS, n=8.

| Group | TNCC<br>(cell/μL) | % | | | | |
| --- | --- | --- | --- | --- | --- | --- |
|  |  | N | L | M | E | MC |
| **sEAS** | 500<br>[257 – 3775] | 83.0<sup>*†</sup><br>[57.0 – 94.0] | 10<sup>*†</sup><br>[1 – 26.5] | 6.5<sup>*†</sup><br>[4.5 – 16.5] | 0 | 0 |
| **mEAS** | 250<br>[150 – 425] | 12.0<sup>†</sup><br>[4.5 – 17.0] | 50<sup>*</sup><br>[34.5 – 56.5] | 36<br>[28.5 – 54.5] | 1.0<br>[0.0 – 5.0] | 0 |
| **Ctr** | 250<br>[200 – 300] | 1.0<sup>*</sup><br>[1.0 – 3.0] | 30<br>[24 – 4] | 68<sup>*</sup><br>[56 – 75] | 0 | 0 |
| **maEAS** | 291.5<br>[169.5 – 355.8] | 2.35<br>[1.18 – 5.03] | 47.65<sup>†</sup><br>[22 – 58.73] | 46<sup>†</sup><br>[28.2 – 70.5] | 0.0<br>[0.0 – 0.7] | 5.0<br>[4.18 – 5.75] |

\*sEAS= Severe Asthma, mEAS= Mild/Moderate Asthma, maEAS= Mastocytic Asthma, Ctr= Healthy; TNCC= Total Nucleated Cell Count (Average recorded); N= neutrophil, L= lymphocyte, M= macrophage, E= eosinophil, MC= mast cell. Note: Medians in the same column with the same superscript symbol differ significantly from each other.

### Mast cell protease gene expression in BALF (UT & NCSU-CVM)

In order to determine whether mast cells have altered protease phenotypes in equine asthma, gene expression analysis was performed on 37 horses (see **Fig. 1** and **Supplementary Table S4 online**). Samples from five horses (4 sEAS, 1 mEAS) were excluded due to poor quality and/or concentrated RNA. Analysis of protease gene expression in BALF isolated cells revealed that chymase was significantly lower in horses with maEAS (fold change: 0.18 [0.08 – 0.54]) than control (1.88 [0.33 – 2.46]; p < 0.05), or sEAS horses (0.92 [0.65 – 1.97]; p < 0.05), implying that mast cell protease expression is altered in horses with mastocytic inflammation. As expected, CD117(ckit), expression was significantly increased in horses with maEAS (fold change: 2.32, [1.31 – 3.85]) compared to sEAS (0.38 [0.27 – 1.50]; p < 0.05). Tryptase expression appeared slightly higher in horses with maEAS compared to sEAS, but was not statistically different between any groups. Additionally, carboxypeptidase A3 was not amplified in any sample, with either primer set identified, but whether this was due to a technical issue or lack of carboxypeptidase A3 gene expression has not been determined.

**Figure 1.**
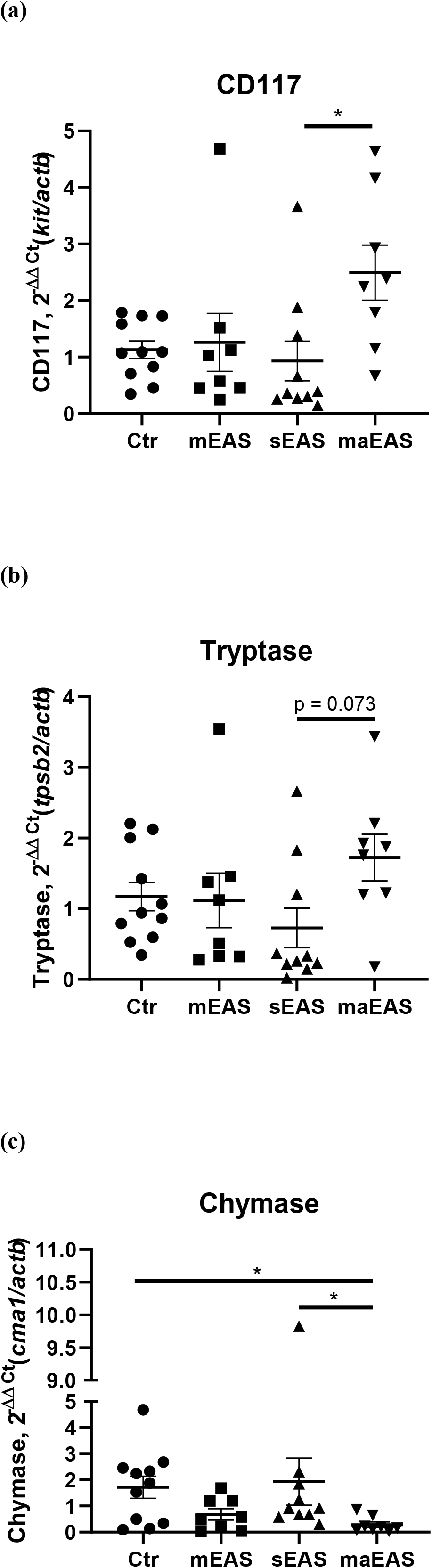
Relative gene expression level of (a) CD117 (*ckit*), (b) tryptase (*tpsb2*), and (c) chymase (*cma1*) in BALF cell pellets in healthy (n = 11), mild/moderate asthma (n = 8), severe asthma (n = 10), and mastocytic asthma (n = 8). All target genes were normalized to the house keeping gene β-actin (*actb*). Displayed figures represent the mean with SEM. sEAS= Severe Asthma, mEAS= Mild/Moderate Asthma, Ctr= Healthy, maEAS= Mastocytic Asthma. *, p<0.05.

### Multiplex Bead Immunoassay Analysis (Interleukin [IL]-2, IL-4, IL-5, IL-17A, CXC motif chemokine ligand [CXCL]8, Interferon [IFN]γ, Tumor Necrosis Factor [TNF]α) (UT-CVM)

Evaluation of BALF cytokine concentrations was only performed in the UT-CVM study population for consistency of the BALF procedure (see **Table 3, Fig. 2, and Supplementary Table S5 online**). Significant differences between groups were found for TNFα, CXCL8 (formerly known as IL-8), and IFNγ. BALF multiplex showed that horses with asthma had significantly higher TNFα (mEAS: p < 0.05; sEAS: p < 0.001) concentrations compared to healthy horses as well as significantly elevated CXCL8 (mEAS: p < 0.05; sEAS: p<0.0001), concentrations compared to healthy horses. Additionally, IFNγ was significantly elevated in sEAS compared to healthy horses (p < 0.05). There were no statistical differences in cytokine concentrations between horses with mEAS or sEAS. No significant differences between groups were found for the remaining analytes.

**Table 3.** Median cytokine concentrations of IL-5, IL-17A, IL-2, IL-4, IFNɣ, CXCL8, and TNFα measured via multiplex bead immunoassay in healthy (n=11), mild/moderate asthma (n = 9), and severe asthma (n = 14). sEAS= Severe Asthma, mEAS= Mild/Moderate Asthma, Ctr= Healthy. Data are expressed as median [IQR].

| Group | IL-5 pg/mL | IL-17A<br>pg/mL | IL-2 pg/mL | IL-4<br>pg/mL | IFNγ<br>pg/mL | CXCL8<br>pg/mL | TNFα pg/mL |
| --- | --- | --- | --- | --- | --- | --- | --- |
| sEAS | 0.99<br>[0.5 – 1.98] | Not<br>detected | 0.46<br>[0.44 – 0.63] | 16.35<br>[16.35 – 16.35] | 35.28*<br>[21.08 – 75.45] | 19.83*<br>[16.86 – 27.82] | 11.31*<br>[3.67 – 21.77] |
| mEAS | 1.05<br>[0.72 – 3.95] | Not<br>detected | 0.44<br>[0.44 – 1.32] | 16.35<br>[16.35 – 24.16] | 28.87<br>[21.08 – 77.64] | 14.41<sup>†</sup><br>[8.74 – 22.48] | 6.54<sup>†</sup><br>[4.79 – 7.69] |
| Ctrl | 0.5<br>[0.5 – 0.99] | Not<br>detected | 0.44<br>[0.44 – 0.44] | 16.35<br>[16.35 – 16.35] | 21.08*<br>[21.08 – 21.08] | 4.54*<sup>†</sup><br>[1.26 – 8.88] | 2.0*<sup>†</sup><br>[0.8 – 3.63] |

Note: Medians in the same column with the same superscript symbol differ significantly from each other.

**Figure 2.**
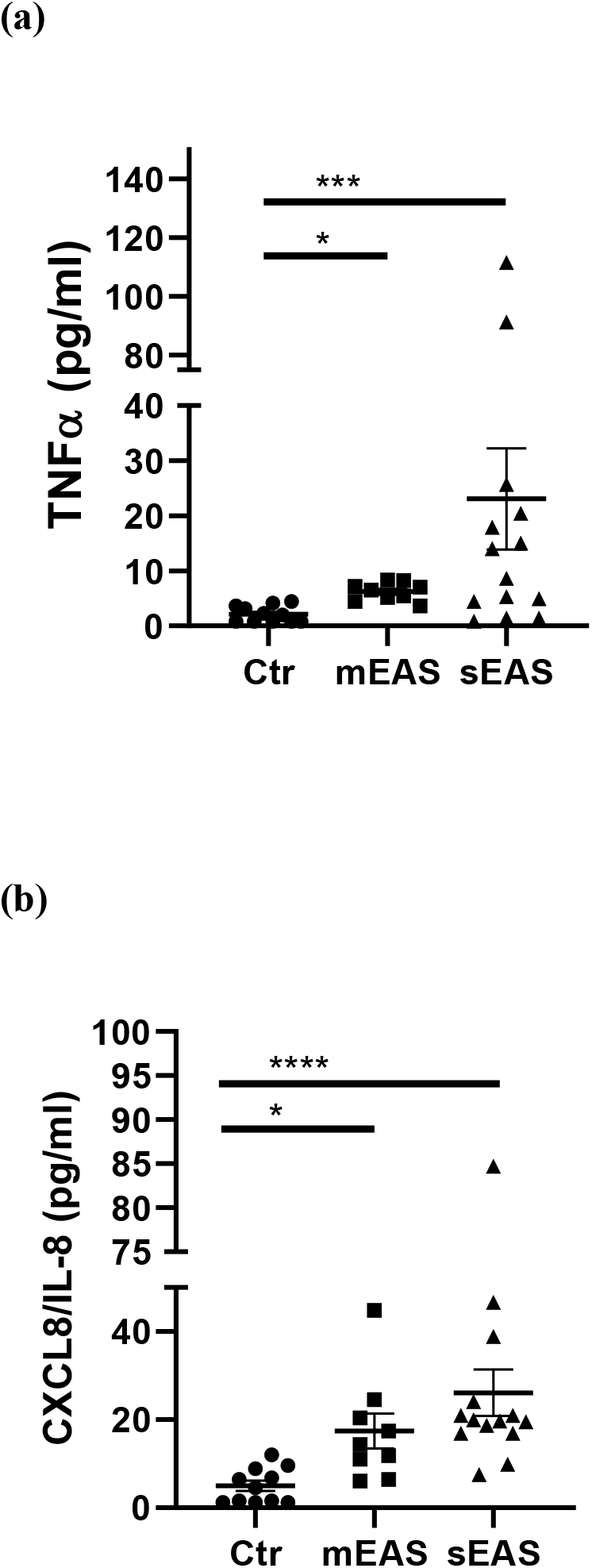
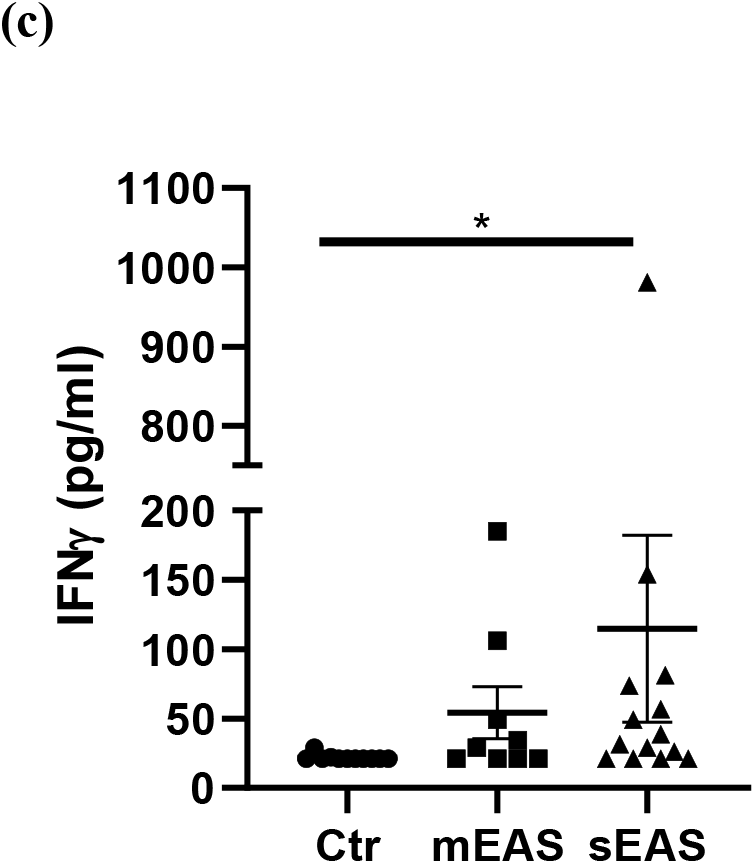
Cytokine concentrations of (a) TNFα, (b) CXCL8/IL-8, and (c) IFNɣ measured via multiplex bead immunoassay in healthy (n=11), mild/moderate asthma (n = 9), and severe asthma (n = 14). Displayed figures represent the mean with SEM. sEAS= Severe Asthma, mEAS= Mild/Moderate Asthma, Ctr= Healthy. *, p<0.05; **, p<0.01; ***, p<.001; **** p<0.0001.

Spike and recovery was lowest for IL-17A and high at low concentrations for IL-4 and TNFα (see **Supplementary Table S6)**. Dilutional linearity was achieved for all analytes (see **Supplementary Fig. S1)**.

### Analysis of Similarity Between Samples (UT-CVM)

Pairwise correlation analysis of the variables demonstrated a moderate positive correlation between neutrophil percentage and CXCL8, TNFα, and IFNγ concentrations (Pearson’s r = 0.54, 0.47, 0.35, respectively) (see **Fig. 3**). The gene expression fold change of CD117(ckit) was positively correlated with tryptase (r = 0.94).

**Figure 3.**
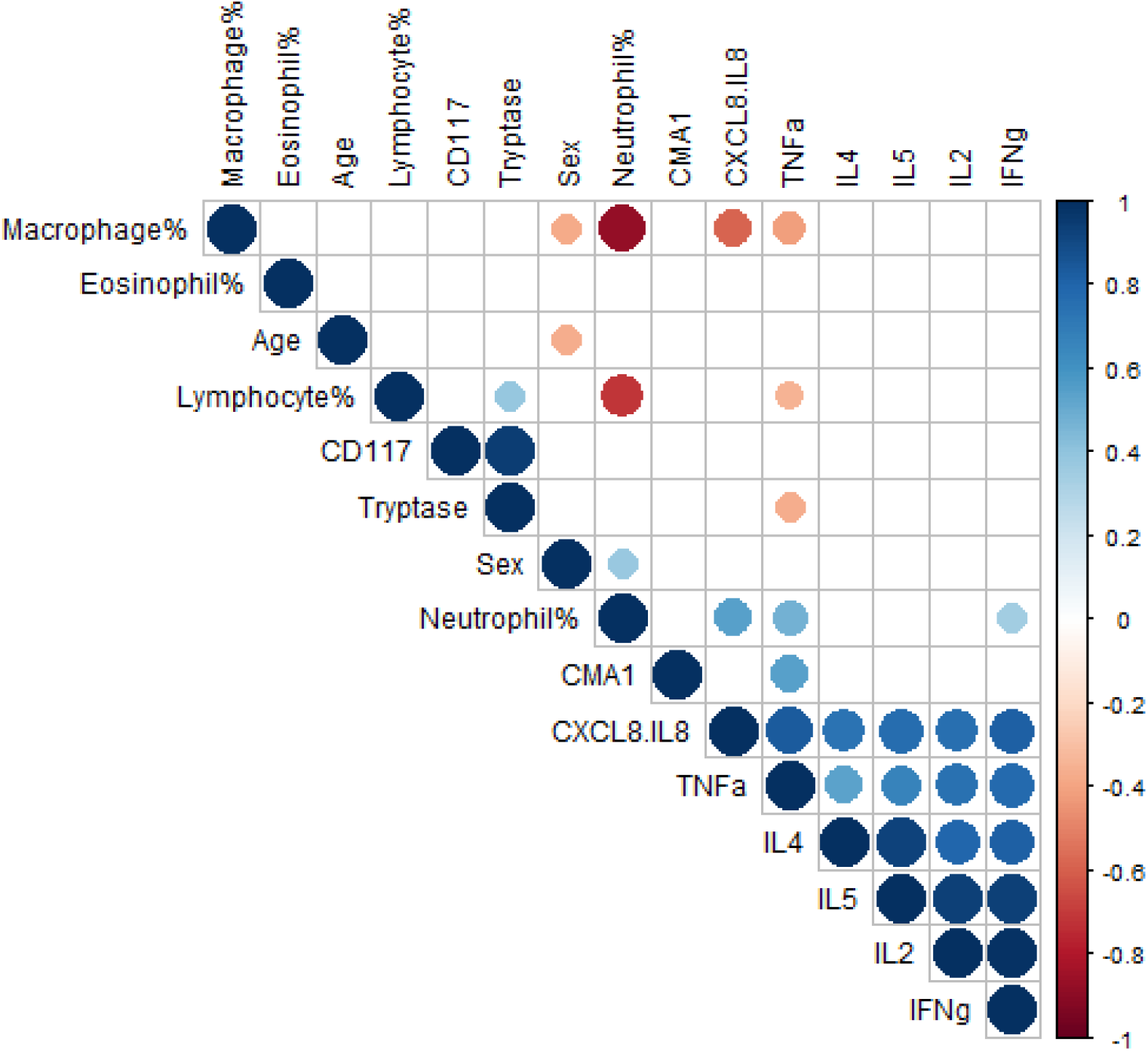
Pairwise correlation of variable. Only significant (p <0.05) correlations are shown. The size of the circle represents the strength of the correlation, and the color represents directionality. The variables are hierarchically clustered based on their correlations.

Multidimensional scaling (MDS) analysis showed separation between samples from horses with sEAS, mEAS, and healthy horses (see **Fig. 4A**). The healthy samples and mEAS samples did not form distinct clusters, but a gradual trend from healthy to mEAS to sEAS was observed on both axes of the two-dimensional projection. Further principal component analysis on samples with complete data showed that the first two principal components (PCs) explained over 98.5% of the variance, with the first PC explaining over 95% of the variance. The absolute value of the loadings of the first two PCs quantified the contribution of each variable to the observed clustering pattern. By ranking the variables on the maximum absolute loadings from PC1 and PC2, IFNγ concentration and neutrophil percentage were the main drivers of the observed clusters (see **Fig. 4B**).

**Figure 4.**
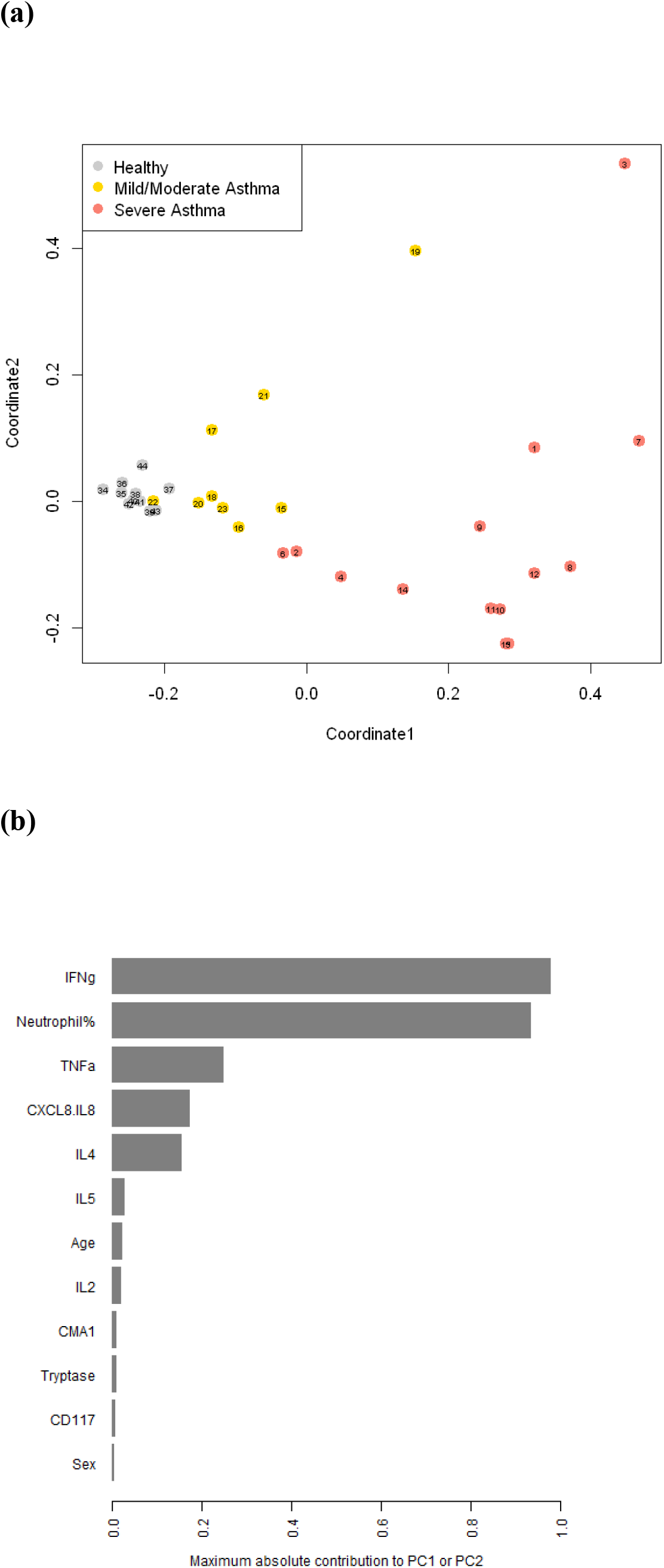
(a) Multidimensional scaling of the samples based on age, sex, cell percentages, gene expression fold changes, and cytokine concentrations. Each sample is labeled with its anonymous numeric identifier. Asthma status is indicated by gray scale and symbol. (b) The maximum contribution to first two principal components from each variable, which shows the relative importance of each variable in driving the observed clustering pattern. The first two principal components explained over 98.5% of the variance in the data.

### Analysis of BMP7 and Tgfβ1 expression (UT & NCSU-CVM)

Potential equine airway remodeling cytokines, BMP7 and Tgfβ1, were evaluated in BALF isolated cells. BMP7 expression in BALF derived cells was not significantly different across the groups, although a subset of horses diagnosed with sEAS had markedly elevated BMP7 expression (see **Fig. 5** and **Supplementary Table S4**). Tgfβ1 was significantly elevated in mEAS (fold change: 1.54, [1.16 – 1.73,] p=0.02) versus healthy (fold change: 1.06, [0.81 – 1.22]). Tgfβ1 to BMP7 ratios did not explain the subgrouping of sEAS seen in BMP7 expression results, although 3 horses in the sEAS group (1 in low BMP7, 2 in high BMP7) were unable to have Tgfβ1 analyzed due to lack of remaining RNA for analysis.

**Figure 5.**
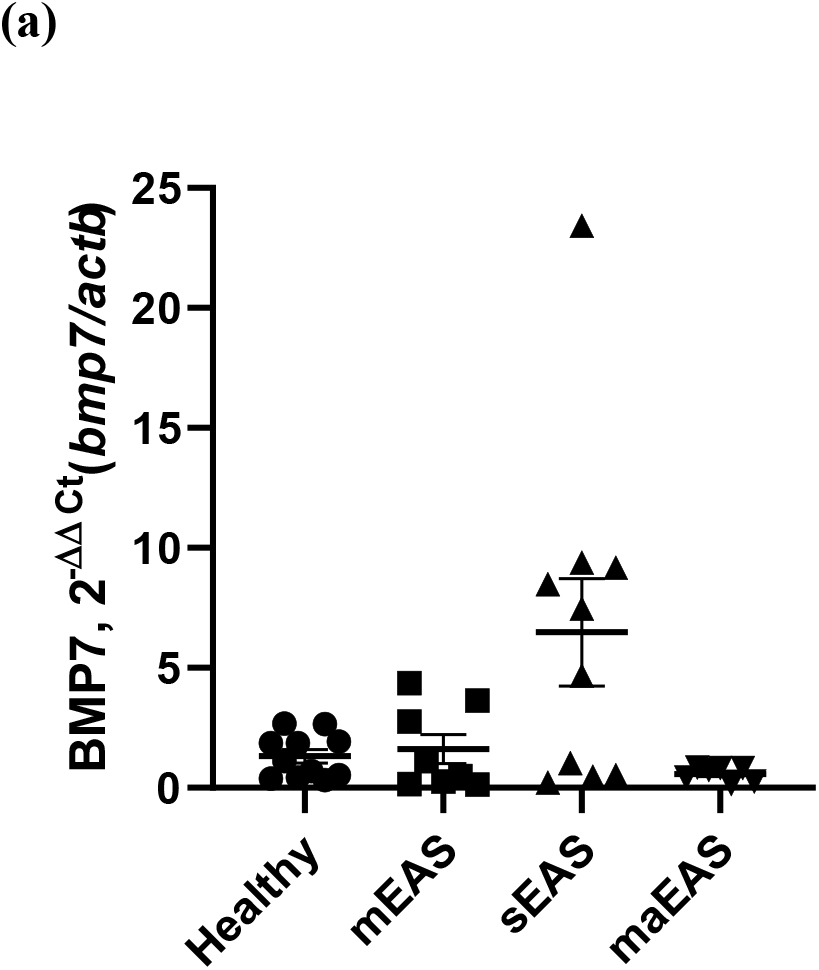
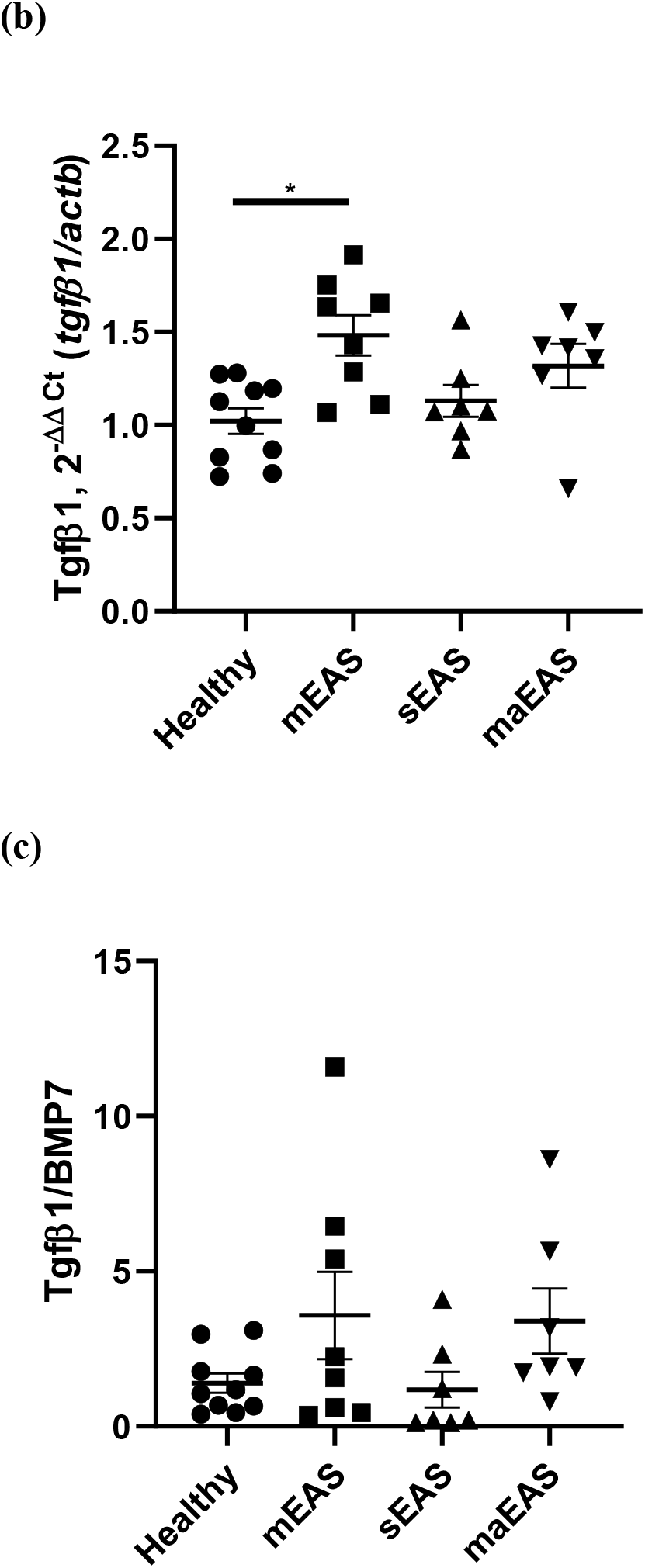
Relative gene expression level of (a) *bmp7*, (b) *tgfβ1*, (c) *tgfβ1/ bmp7* in BALF cell pellets. Sample size of (a) and (b), healthy: n=11, mild/moderate asthma: n = 8, severe asthma: n = 10 (BMP7 low = 4 & BMP7 high = 6), and mastocytic asthma: n = 7. Sample size of (c) and (d), healthy: n=10, mild/moderate asthma: n = 8, severe asthma: n = 7, and mastocytic asthma: n = 7. sEAS= Severe Asthma, mEAS= Mild/Moderate Asthma, Ctr= Healthy, maEAS= Mastocytic Asthma. *, p<0.05; **, p<0.01.

## Discussion

This study, to our knowledge, is the first to examine BALF supernatant cytokine/chemokine concentrations at the protein level, and mast cell protease expression in cells isolated from BALF in horses with asthma. We utilized an unbiased approach and multidimensional scaling to integrate clinical data, cytokine and chemokine concentrations, and mast cell protease expression, in order to investigate the most important biomarkers of equine asthma. This analysis indicated that IFNγ is a key biomarker of equine asthma, in particular, severe equine asthma. Similarly, in BALF cells isolated from humans with severe asthma, IFNγ expression was increased compared to mild/moderate asthma, and further mechanistic analysis of the IFNγ pathway in a mouse model demonstrated that IFNγ dysregulation was the main cause of airway hyper-responsiveness via regulation of the epithelial cell-expressed protein *SLPI* (Secretory Leukocyte Peptidase Inhibitor), which downregulates airway reactivity [33]. In addition to airway hyper-responsiveness, airway remodeling is another key feature of asthma. Tgfβ1 and BMP7 expression in asthma models and human asthma highlights their balance as an important factor in asthma airway remodeling [18–20]. This study is the first to evaluate BMP7 expression in the airway of the horse. Our results demonstrated that a subset of horses with sEAS had elevated BMP7 expression compared to all other groups. BMP7 is a cytokine in the TGFβ superfamily that has both anti-inflammatory and anti-fibrotic effects. The potential role of BMP7 in human asthma has been suggested to be anti-inflammatory, with BMP7’s ability to block detrimental Tgfβ1effects, such as fibroblast activation with subsequent airway remodeling, production of extracellular matrix, and collagen deposition [19, 20]. The sEAS BMP7-high group may have an over exuberant BMP7 response in attempt to counteract airway remodeling. Airway function tests may be worse in the sEAS BMP7 high subset of horses and have less reversible airway obstruction. Further studies to define the horses with BMP7-high and BMP-7 low sEAS may elucidate further subgroupings of sEAS and further define the pathophysiology.

In this study, cytokine/chemokine concentrations in BALF in the main study population (UT-CVM), revealed elevated TNFα and CXCL8 in both mEAS and sEAS horses; therefore, these are potential targets for treatment in asthma. Elevated TNFα and CXCL8 parallels findings in human asthma, in which asthmatics had increased TNFα and CXCL8 versus control, and neutrophilic asthma was associated with further increased CXCL8 [34, 35]. Additionally, sEAS clustered separately from the other groups based on increased IFNγ and neutrophil percentage, therefore they appear to be the best targets for modulation during exacerbation of severe asthma.

We additionally demonstrated that mast cell protease expression is altered in horses with mastocytic asthma, in particular, that chymase is reduced. Loss of chymase may contribute to disease progression since chymase has anti-inflammatory activities in allergic asthma models via modulating IL-33 concentrations, and may be protective of lung function [36, 37]. In contrast, mast cell protease expression is relatively unchanged in mEAS and sEAS. We also demonstrated that mast cells in equine BALF express both tryptase and chymase mRNA. Whether equine mast cells also express carboxypeptidase A3 remains to be determined.

Previous work evaluating mast cell types in horses have investigated lung tissue samples, but not BALF, and reports have been conflicting. A report comparing types in sEAS versus control after moldy hay challenge found no MC_TC_, although severely asthmatic horses had significantly more MC_C_ in the wall of the bronchi [31]. In conflict to this report, another study found increased tryptase expression in lung tissue from asthmatic horses versus control [32]. BALF IgE-bearing cells have been evaluated in asthmatic horses and were not different from control horses, although this is not mast cell-specific and the exact cell type bearing the IgE was not determined [38]. The current study suggests protease dysregulation, and changes in mast cell type, occur in horses with mastocytic asthma.

Both human mast cell types, MC_T_ and MC_TC_, are noted to be increased in the lungs of asthmatics compared to control subjects’ lungs. Additionally, increased numbers of MC_T_ in smooth muscle of subjects with asthma have been associated with airway hyper-reactivity [25, 39]. Corticosteroid use specifically decrease MC_T_ numbers and an increase in MC_TC_ type may be protective of lung function in severe, steroid-dependent human asthmatics [25, 37]. Additional molecular analysis of lung samples revealed that tryptase and carboxypeptidase A3 were among the most differentially expressed genes in asthmatics versus control, with Th2-high asthma associated with expression of tryptase and carboxypeptidase A3, but not chymase [40, 41]. This all suggests the importance of mast cell phenotype in asthma type, pathogenesis, and response to therapy.

Primary limitations of the study are small sample size, which limits extrapolation of results to other areas of the country, in addition to disease and group classification. Definitions of EAS are continuously changing, with current emphasis placed on clinical signs, BALF cytological examination, and ruling out other causes of lower airway inflammation. Tryptase and chymase are primarily made by mast cells, and although mast cell protease knockout models have suggested a mild decrease in basophil number, suggesting low protease expression; basophils are not noted to be in equine BALF [42–45]. In our study, CD117/c-kit expression increased in maEAS horses and highly correlated with tryptase expression levels, therefore the tryptase gene expression is likely to be of mast cell origin, since basophils do not retain CD117 expression once differentiated [46, 47]. CD117 can also be expressed by other immune cell types, specifically type 2 innate lymphoid cells, however these cells have not yet been described in the horse [48]. It is unclear whether the negative CPA3 amplification in all horses in this study is due to a technical problem with the assay or whether horse BALF mast cells truly do not express CPA3. Carboxypeptidase A3 expression in horses warrants further investigation.

Additionally, there is the possibility that the variations in the method of BALF collection, such as the location sampled or volume of return could have influenced cytokine results. Unfortunately, there is not a universally accepted way to normalize the cytokine/chemokine concentrations obtained in BALF recovered across patients. To mitigate this variability, we performed BALF collections in a standardized fashion with standardized fluid volumes infused. Normalization of analyte concentration to fluid volume recovered during the BAL procedure has been recommended, but is not consistently employed in the literature [49].

We propose that further, larger sample size evaluation of protease and cytokine/chemokine profiles will identify biomarkers that support defining equine asthma subtypes based on BALF cellular infiltrate, including eosinophilic mild/moderate asthma (emEAS), mastocytic mild/moderate asthma (maEAS), and severe asthma (sEAS). These groups likely have differences that will require specific environmental management and treatments for optimal management of the patient. Lung function tests would be additional useful information for support of specific treatments. A staging scale for severe asthma could be used in addition to, or substitution of, lung function tests, but a staging scale for mEAS does not exist at this time [50].

In conclusion, this study demonstrates that, similar to human patients with asthma, horses with mEAS or sEAS have increases in CXCL8 and TNFα, with IFNγ additionally separating sEAS from other groups. In addition, as seen in humans with Th2-high asthma, maEAS is associated with an increase in tryptase and decrease in chymase. Therefore, horses may serve as a naturally-occurring model of human asthma and potential model for chymase dysregulation.

## Methods

### Main study population – (UT-CVM)

Horses were enrolled from the institution’s equine hospital caseload and research herd from February 2018 to April 2019. The procedures were approved by the University of Tennessee Institutional Animal Care and Use Committee (protocol 2533). Horses from the clinical caseload were enrolled on a volunteer basis with written, informed owner consent, and completed standardized questionnaire.

**Table 1** contains the diagnostic criteria for each experimental group, in part defined using consensus guidelines [1]. Attending large animal internal medicine clinicians at the time of client-owned horse enrollment evaluated the horses. The institution’s research herd was evaluated by the same two investigators for all horses (JW and CS). Exclusion criteria included horses less than one year of age, history of systemic or respiratory disease (besides suspected asthma) in the past three months, and corticosteroid (systemic or intranasal) administration begun or significant dosage changes within two weeks of enrollment. All horses underwent evaluation via history and physical exam, including a rebreathing exam, CBC/fibrinogen measurement, and BALF cytology. Horses were excluded from analysis if they did not meet final inclusion criteria.

### Sub-population – (NCSU-CVM)

As data collection progressed and BALF results of the UT-CVM horses were evaluated, it was observed that no horses with ≥ 1% mast cells were enrolled. Therefore, additional equine BALF samples were obtained from a collaborator (KS) at NCSU-CVM, who had previously identified horses with mastocytic asthma (maEAS). These additional samples were used for investigation of mast cell protease mRNA expression because this parameter could be normalized by using standardized quantities of cDNA. The NCSU-CVM samples were processed separately from the remainder of the samples and are identified as a sub-population (**see Table 1**).

### Sample collection

#### Main Population (UT-CVM)

Blood was collected from the jugular vein into K_2_EDTA tubes (Becton, Dickson, and Co., Franklin Lakes, NJ) prior to sedation and was submitted to the institution’s clinical pathology laboratory for CBC determination using an Advia 2120i hematology instrument (Siemens Healthcare Diagnostics, Inc.m Tarrytown, NY) and heat-precipitated fibrinogen measurement using an analogue Goldberg TS meter clinical refractometer (Reichert Technologies, Buffalo, NY) by a licensed medical technologist.

Bronchoalveolar lavage procedure was performed on standing, sedated horses using a blind technique with a BAL catheter (Mila International, Florence, KY). Sedation for each horse was determined by the attending clinician and consisted of xylazine (0.2 – 0.5 mg/kg) or detomidine (5 - 10 μg/kg), combined with butorphanol (10 – 20 μg/kg) IV. The BAL catheter was passed nasotracheally until wedged. Once wedged, the cuff was inflated and 200 mL of sterile saline was infused and re-aspirated by 60 mL syringe. The first 10 mL of aspirated sample was discarded, with subsequent BALF collected and pooled for analysis.

#### Sub-Population (NCSU-CVM)

BAL fluid was collected in a similar manner, with the exception of a larger infusion volume. Following sedations with detomidine (0.005 – 0.01 mg/kg IV) and butorphanol (0.02 – 0.04 mg/kg IV), a cuffed catheter (Bivona, outside diameter 11 mm, 244 cm length, Smith Medical) was passed nasotracheally until wedged. The cuff was then inflated and 300 mL of warmed sterile saline solution was infused and re-aspirated by 60 mL syringe, 150 mLs at a time. The aspirated fluid was pooled, placed on ice and processed within 30 minutes of collection. Of the recovered fluid, 10 mL was submitted to NCSU’s Clinical Pathology Service for a total nucleated cell count and 300 cell differential cell count performed by blinded clinical pathologists. Slides were prepared within 2 hours of sample collection, and both direct smear and cytocentrifuge slides were examined. The remaining fluid was filtered to remove mucous and centrifuged to collect the cell pellet. Aliquots were stored at −80°C until further analysis.

### BALF Cell Count and Cytologic Analysis (UT-CVM)

BALF clinicopathologic analysis consisted of a TNCC and cytologic examination of cytocentrifuged BALF. To optimize preservation of cell morphology, BALF was placed in K_2_EDTA tubes for TNCC and cytologic evaluation.

TNCC was performed using a scil Vet ABC hematology analyzer (scil animal care, Gurnee, IL) by a trained laboratory assistant or medical technologist. The WBC reported by the instrument was reported as TNCC. To prepare BALF for TNCC, 200 μL of well-mixed BALF was placed into a previously prepared hyaluronidase-containing cryo tubes, in order to break down any mucus that could clog the instrument. All TNCC measurements were done in duplicate to mitigate measurement imprecision, and average of the duplicate TNCC was reported.

For cytologic evaluation, 100μL of well-mixed BALF was placed in a cytocentrifuge (Aerospray 7120 Slide Stainer and Cytocentrifuge, Wescor Incorporated, Logan UT). Cytocentrifuged specimens were air-dried and stained with aqueous Wright’s stain. Nucleated cell differential counting (minimum of 100 cells) was performed by the pathologist on duty and reported as cell type percentages; respiratory epithelial cells were excluded from these differential counts.

### BALF Preparation for RNA and Cytokine Analysis (UT-CVM)

Pooled BALF placed in non-K_2_EDTA tubes was refrigerated at 4°C until further processing, within two hours of collection. To prepare specimens for cytokine and RNA analysis, BALF was strained through sterile 4×4 gauze, followed by a 70 μm nylon cell strainer to remove mucus, and then centrifuged. Aliquots of supernatant were made and stored at −80°C until further analysis. Cell pellets were washed two additional times and were stored dry at −80°C until further processing.

### RNA extraction and gene expression (UT and NCSU-CVM)

For the main population, RNA was extracted from BALF cell pellets using RNeasy Plus Mini Kit (Qiagen, Hilden, Germany) per manufacturer instructions. For the sub-population, RNA was isolated using an RNeasy Mini Kit with the addition of RNase-free DNase kit (Qiagen) per manufacturer’s instructions. The isolated RNA samples were stored at −80°C for up to 18 months.

For both populations, quantification and purity of RNA extracted was assessed via NanoDrop 2000c Spectrophotometer (Thermo Fisher Scientific, Waltham, MA). RNA integrity was analyzed on a subset of samples using 2100 Bioanalyzer and the RNA 6000 Nano kit (Agilent, Santa Clara, CA). Reverse transcription and cDNA synthesis was performed on 200ng of RNA using Maxima First Strand cDNA synthesis kit (Thermo Fisher Scientific) and stored at −80°C. Taqman Gene Expression Assays were used for all target genes; β-actin, CD117 (or ckit), tryptase, chymase (CMA1), carboxypeptidase A3 (CPA3), Tgfβ1, & BMP7 (Applied Biosystems, Foster City, CA) (see **Supplementary Table S7**). Real time PCR was performed on 10ng cDNA, with Taqman Fast Advanced Master Mix (Applied Biosystems). QuantStudio 6 Flex Real-Time PCR system (Applied Biosystems) was used for thermal cycling at recommended settings. Results were normalized to β-actin and relative quantification was performed using 2^−ΔΔCt^ method. Results from plates run were used when β-actin standard deviations were ≤ 0.7. Average standard deviation of all plates run was 0.55.

### Multiplex Bead Immunoassay Analysis (UT-CVM)

Previously collected BALF supernatant was thawed at room temperature, then hyper-centrifuged to remove any precipitates formed during thawing. Cytokines/chemokines included in the assay were: IL-2, IL-4, IL-5, IL-17A, CXCL8, IFNγ, and TNFα. Cytokine concentrations were quantified using an equine-specific Milliplex® Map Magnetic Bead Panel (EMD Millipore, St Louis, MO, USA) according to the manufacturer’s instructions using a Luminex® 200 instrument and Luminex xPONENT® software run in duplicate (Luminex, Austin, TX, USA). Dilutional linearity and percent recovered were determined using low-cytokine containing BALF spiked with kit standard.

Data analysis was done using Milliplex Analyst v5.1 software (EMD Millipore). A bead count of at least 50 beads per well was used for inclusion in analysis and individual samples with a coefficient of variation (CV) > 15% were not included. The Analyst® software assigned the lowest detectable concentration for the individual analytes in which the cytokine/chemokine concentrations fell below the lower limit of detection (LLOD) of the assay. Best fit standard curve for each analyte was used to calculate analyte concentrations.

### Statistical analysis

Normality of data was assessed using Shapiro-Wilk test. Descriptive data are expressed as median with interquartile range for both normal and non-normally distributed data for consistency. Descriptive data, protease expression, and cytokine concentrations were analyzed using one-way analysis of variance with Tukey’s multiple comparisons in normally distributed data, and by Kruskall-Wallis with Dunn’s multiple comparisons in non-normally distributed data. Pairwise correlation of variables including age, sex, cell percentages, gene expression fold changes, and cytokine concentrations was performed, with additional multidimensional scaling (MDS) analysis in order to visualize on a two-dimensional figure the clustering of samples based on age, sex, gene expression fold changes, cytokine concentrations, and neutrophil percentage. Since cell percentages add up to 100, the percentage of only one cell type was used in the clustering to avoid spurious signal driven by cell percentages going in opposite directions. Bray-Curtis distance function was applied to calculate the dissimilarity matrix of dataset, which ignored missing gene expression values in 5 samples. To evaluate which variable contribute the most to sample clustering, an additional principal components analysis (PCA, which is a specific solution of MDS based on Euclidean distance of complete data) on the samples with no missing values was performed, followed by quantification of the contributions of each variable to the first two principal components. Significance was set at p ≤ 0.05.

## Acknowledgements

This work was funded by the Morris Animal Foundation (D18EQ-830) and Boehringer Ingelheim Vetmedica, Inc. (Advancement in Equine Research Award). Dr. Lennon was supported by a U.S. National Institutes of Health Mentored Research Scientist Development Award (K01 OD019729).

## Author Contributions

J.S.W. and E.M.L. conceived the project. J.S.W., M.H., C.S., B.F., and E.M.L. planned the study design. J.S.W. collected and processed samples, performed gene expression and cytokine analysis, analyzed data, and wrote the first draft of the manuscript. B.F. optimized TNCC procedures and provided guidance on clinicopathologic data. M.H., C.S., M.H., and J.S.W. collected UT-CVM samples, and K.D. and M.K.S. collected NCSU samples and extracted RNA. J.S.W. processed all UT-CVM samples. M.H., C.S., B.F., and E.M.L. edited grant proposals for the support of this research. Y.L. and P.W. performed advanced statistical analysis and associated manuscript figures. J.S.W. and E.M.L. created all other manuscript figures. E.M.L. supervised J.S.W. and edited all drafts of the manuscript. M.H., C.S., B.F., K.D., M.K.S., Y.L, and P.W. edited manuscript. All the authors have read and approved the final manuscript.

## Additional Information

The author(s) declare no competing interests.

