## Supplemental Information for "Multidimensional Analysis of Bronchoalveolar Lavage Cytokines and Mast Cell Proteases Reveals Interferon-γ as a Key Biomarker in Equine Asthma Syndrome"

Jane S. Woodrow, DVM, MS, DACVIM^1,2^; Melissa Hines, DVM, PhD, DACVIM^3^; Carla Sommardahl, DVM, PhD, DACVIM^3^; Bente Flatland, DVM, MS, DACVP, DACVIM^4^; Kaori U. Davis, DVM^5^; Yancy Lo, PhD^6^; Zhiping Wang, PhD^6^; Mary Katherine Sheats, DVM, PhD, DACVIM^5;^ Elizabeth M. Lennon, DVM, PhD, DACVIM^2^

**Supplemental Tables and Figures**

**Supplementary Table S1.** Experimental groups: history & clinical signs, rebreathing exam, sex, age, & breed for individual horses in each group.

|  | **Diagnosis** | **Clinical Signs** | **Rebreathing Exam** | **Sex** | **Age (yrs)** | **Breed** |
| --- | --- | --- | --- | --- | --- | --- |
| Horse 1 | sEAS | Chronic cough | Delayed recovery | M | 20 | Pony |
| Horse 2 | sEAS | Chronic cough, intermittent nasal discharge | NSF | M | 13 | Not reported |
| Horse 3 | sEAS | Chronic cough | Tracheal rattle | M | 13 | MFT |
| Horse 4 | sEAS | Chronic cough, intermittent nasal discharge | Crackles, wheezes | M | 13 | ASH |
| Horse 5 | sEAS | Chronic cough | Crackles, wheezes | G | 12 | Pony |
| Horse 6 | sEAS | Increased respiratory effort | Wheezes | M | 14 | TWH |
| Horse 7 | sEAS | Chronic cough, exercise intolerance | Wheezes | G | 19 | Not reported |
| Horse 8 | sEAS | Chronic cough | NSF | G | 20 | Arab |
| Horse 9 | sEAS | Increased respiratory rate at rest | Wheezes | G | 14 | TWH |
| Horse 10 | sEAS | Chronic cough, intermittent nasal discharge | Crackles, wheezes | M | 20 | Arab |
| Horse 11 | sEAS | Wheezes and increased respiratory effort at rest | Wheezes | G | 10 | TWH |
| Horse 12 | sEAS | Chronic intermittent cough and increased respiratory rate | Wheezes, occasional crackle, increased respiratory effort | G | 19 | H |
| Horse 13 | sEAS | Chronic intermittent cough | Wheezes and worsened cough | G | 12 | QH |
| Horse 14 | sEAS | Cough, increased respiratory effort | Worsened respiratory effort and cough | G | 17 | QH |
| Horse 15 | mEAS | Increased respiratory rate in barn | NSF | M | 18 | QH |
| Horse 16 | mEAS | Cough, intermittent nasal discharge | Cough upon tracheal palpation, delayed recovery | G | 13 | QH |
| Horse 17 | mEAS | None observed | Delayed recovery | M | 22 | QH |
| Horse 18 | mEAS | Cough when in barn | Delayed recovery | M | 21 | QH |
| Horse 19 | mEAS | Chronic cough | Wheezes | G | 14 | MFT |
| Horse 20 | mEAS | Increased respiratory rate, exercise intolerance | NSF | G | 14 | ASH |
| Horse 21 | mEAS | Exercise intolerance, intermittent nasal discharge | Delayed recovery, nasal discharge | G | 11 | PF |
| Horse 22 | mEAS | Chronic cough | Crackles | G | 12 | TWH |
| Horse 23 | mEAS | Chronic cough | Wheezes | M | 18 | QH |
| Horse 24 | Ctr | None | NSF | M | 8 | TWH |
| Horse 25 | Ctr | None | NSF | M | 18 | QH |
| Horse 26 | Ctr | None | NSF | M | 19 | QH |
| Horse 27 | Ctr | None | NSF | M | 16 | QH |
| Horse 28 | Ctr | None | NSF | M | 14 | MFT |
| Horse 29 | Ctr | None | NSF | M | 17 | QH |
| Horse 30 | Ctr | None | NSF | M | 26 | APH |
| Horse 31 | Ctr | None | NSF | M | 16 | TWH |
| Horse 32 | Ctr | None | NSF | G | 11 | TWH |
| Horse 33 | Ctr | None | NSF | M | 14 | QH |
| Horse 34 | Ctr | None | NSF | G | 12 | TB |
| Horse 35 | maEAS | None | NSF | M | 6 | QH |
| Horse 36 | maEAS | Abdominal lift | NSF | M | 18 | QH |
| Horse 37 | maEAS | None | NSF | M | unknown | unknown |
| Horse 38 | maEAS | Abdominal lift, nasal discharge | NSF | M | unknown | unknown |
| Horse 39 | maEAS | None | NSF | G | 9 | TBx |
| Horse 40 | maEAS | Nostril flare | Delayed recovery | M | unknown | unknown |
| Horse 41 | maEAS | None | NSF | M | 3 | unknown |
| Horse 42 | maEAS | Cough, abdominal lift | NSF | M | 3 | unknown |

*sEAS= Severe Asthma, mEAS= Mild/Moderate Asthma, Ctr= Healthy, maEAS= Mastocytic Asthma; M= Mare, G= Gelding; NSF= No significant findings; MFT= Missouri Fox Trotter, ASH= American Saddle Horse, TWH= Tennessee Walking Horse, H= Holsteiner, QH= Quarter Horse, PF= Paso Fino, TB= Thoroughbred, APH= American Paint Horse, TBx= Thoroughbred cross.

**Supplementary Table S2.** Bronchoalveolar lavage fluid characteristics: fluid volume recovered, % volume recovered, TNCC, neutrophil %, lymphocyte %, macrophage %, eosinophil %, and mast cell % for individual horses in each group. Data is represented at the median [IQR] for each group.

|  |  |  |  |  |  | **%** |  |  |
| --- | --- | --- | --- | --- | --- | --- | --- | --- |
| **Horse** | **mLs Fluid Recovered** | **% Volume Recovered** | **TNCC**  **(cell/µL)** | **N** | **L** | **M** | **E** | **MC** |
| **sEAS** |  |  |  |  |  |  |  |  |
| 1 | 23 | 11.5 | 13,000 | 68 | 17 | 15 | 0 | 0 |
| 2 | 62 | 31 | 900 | 36 | 31 | 33 | 0 | 0 |
| 3 | 120 | 60 | 49,000 | 98 | 1 | 1 | 0 | 0 |
| 4 | 52 | 26 | 2,000 | 48 | 31 | 21 | 0 | 0 |
| 5 | 24 | 12 | 250 | 94 | 1 | 5 | 0 | 0 |
| 6 | 78 | 39 | 200 | 35 | 25 | 40 | 0 | 0 |
| 7 | 19 | 9.5 | Unable to determine | 97 | 0 | 3 | 0 | 0 |
| 8 | 30 | 15 | 5,550 | 94 | 3 | 3 | 0 | 0 |
| 9 | 58 | 29 | 300 | 67 | 18 | 15 | 0 | 0 |
| 10 | 70 | 35 | 1,150 | 84 | 11 | 5 | 0 | 0 |
| 11 | 40 | 40 | 500 | 82 | 9 | 9 | 0 | 0 |
| 12 | 60 | 60 | 200 | 88 | 6 | 6 | 0 | 0 |
| 13 | 46 | 46 | 300 | 92 | 1 | 7 | 0 | 0 |
| 14 | 81 | 81 | 350 | 60 | 35 | 5 | 0 | 0 |
| ***Median [IQR]*** | 55  [28.5 – 72] | 27.5  [14.25 – 36.0] | 500  [257 – 3775] | 83.0  [57.0 – 94.0] | 10  [1 – 26.5] | 6.5  [4.5 – 16.5] | 0 | 0 |
| **mEAS** |  |  |  |  |  |  |  |  |
| 15 | 70 | 35 | 1,000 | 14 | 69 | 17 | 0 | 0 |
| 16 | 120 | 60 | 400 | 21 | 42 | 36 | 1 | 0 |
| 17 | 90 | 45 | 250 | 1 | 50 | 44 | 5 | 0 |
| 18 | 75 | 37.5 | 150 | 7 | 28 | 59 | 6 | 0 |
| 19 | 65 | 32.5 | 450 | 20 | 54 | 25 | 1 | 0 |
| 20 | 80 | 40 | 150 | 10 | 58 | 32 | 0 | 0 |
| 21 | 100 | 50 | 250 | 12 | 37 | 50 | 1 | 0 |
| 22 | 60 | 30 | 150 | 2 | 32 | 61 | 5 | 0 |
| 23 | 78 | 39 | 300 | 12 | 55 | 33 | 0 | 0 |
| ***Median [IQR]*** | 78  [67.5 – 95] | 39.0  [33.8 – 47.5] | 250  [150 – 425] | 12.0  [4.5 – 17.0] | 50  [34.5 – 56.5] | 36  [28.5 – 54.5] | 1.0  [0.0 – 5.0] | 0 |
| **Ctr** |  |  |  |  |  |  |  |  |
| 24 | 70 | 35 | 200 | 0 | 25 | 75 | 0 | 0 |
| 25 | 120 | 60 | 300 | 1 | 31 | 68 | 0 | 0 |
| 26 | 50 | 25 | 100 | 0 | 44 | 56 | 0 | 0 |
| 27 | 60 | 30 | 400 | 3 | 50 | 47 | 0 | 0 |
| 28 | 60 | 30 | 250 | 1 | 37 | 62 | 0 | 0 |
| 29 | 90 | 45 | 300 | 1 | 13 | 86 | 0 | 0 |
| 30 | 80 | 40 | 200 | 4 | 24 | 72 | 0 | 0 |
| 31 | 70 | 35 | 400 | 1 | 25 | 74 | 0 | 0 |
| 32 | 40 | 16 | 200 | 3 | 15 | 82 | 0 | 0 |
| 33 | 120 | 60 | 300 | 6 | 30 | 64 | 0 | 0 |
| 34 | 88 | 44 | 250 | 3 | 45 | 52 | 0 | 0 |
| ***Median [IQR]*** | 70  [60 – 90] | 35.0  [30.0 – 45.0] | 250  [200 – 300] | 1.0  [1.0 – 3.0] | 30  [24 – 44] | 68  [56 – 75] | 0 | 0 |
| **maEAS** |  |  |  |  |  |  |  |  |
| 35 | 155 | 51.7 | 118 | 1.7 | 40 | 54 | 0.7 | 3.6 |
| 36 | 160 | 53.3 | 145 | 2.7 | 45.3 | 47 | 0 | 5 |
| 37 | 150 | 50 | 300 | 3 | 59.3 | 33 | 0 | 4.7 |
| 38 | 160 | 53.3 | 283 | 5.7 | 62 | 26.6 | 0.7 | 5 |
| 39 | 200 | 66.7 | 313 | 1 | 50 | 45 | 0 | 4 |
| 40 | 110 | 36.7 | 788 | 9 | 57 | 25 | 2 | 7 |
| 41 | 220 | 73.3 | 370 | 0 | 16 | 79 | 0 | 5 |
| 42 | 160 | 53.3 | 243 | 2 | 16 | 76 | 0 | 6 |
| ***Median [IQR]*** | 160  [151.3 – 190] | 53.3  [50.4 – 63.4] | 291.5  [169.5 – 355.8] | 2.35  [1.18 – 5.03] | 47.65  [22 – 58.73] | 46  [28.2 – 70.5] | 0.0  [0.0 – 0.7] | 5.0  [4.18 – 5.75] |

*sEAS= Severe Asthma, mEAS= Mild/Moderate Asthma, maEAS= Mastocytic Asthma, Ctr= Healthy; TNCC= Total Nucleated Cell Count (Average recorded); N= neutrophil, L= lymphocyte, M= macrophage, E= eosinophil, MC= mast cell.

**Supplementary Table S3.** Excluded horses from the study based on history, physical exam, and diagnostics.

| **ID** | **Suspicion based on history & physical exam** | **Reason for exclusion based on diagnostics** |
| --- | --- | --- |
| 1 | Asthma | Elevated lymphocytes on BAL cytology |
| 2 | Asthma | Elevated lymphocytes on BAL cytology |
| 3 | Asthma | Elevated lymphocytes on BAL cytology |
| 4 | Asthma | Normal BAL cytology |
| 5 | Asthma | Normal BAL cytology |
| 6 | Asthma | Normal BAL cytology |
| 7 | Asthma | Elevated lymphocytes on BAL cytology |
| 8 | Asthma | Normal BAL cytology |
| 9 | Asthma | Elevated lymphocytes on BAL cytology |
| 10 | Asthma | Normal BAL cytology |
| 11 | Asthma | BAL cytology unreadable |
| 12 | Asthma | BAL cytology unreadable |
| 13 | Asthma | Additional upper airway issues: DDSP and laryngeal hemiplegia |
| 14 | Asthma | Mature neutrophilia on CBC |
| 15 | Healthy | Elevated lymphocytes on BAL cytology |
| 16 | Healthy | Elevated neutrophils on BAL cytology |
| 17 | Healthy | Elevated neutrophils on BAL cytology |
| 18 | Healthy | Elevated lymphocytes on BAL cytology |
| 19 | Healthy | Elevated lymphocytes on BAL cytology |
| 20 | Healthy | Elevated neutrophils on BAL cytology |

**Supplementary Table S4.** Relative gene expression level of CD117, tryptase, chymase, BMP7, Tgfβ1, and Tgfβ1/BMP7. Data is represented at the median [IQR] for each group.

| **Group** | **CD117** | **Tryptase** | **Chymase** | **BMP7** | **TGFβ1** | **TGFβ1/BMP7** |
| --- | --- | --- | --- | --- | --- | --- |
| **Ctr** | 1.09  [0.70 – 1.73] | 0.94  [0.60 – 2.01] | 1.88  [0.33 – 2.46] | 1.12  [0.42 – 1.93] | 1.06  [0.81 – 1.22] | 1.12  [0.61 – 2.07] |
| **mEAS** | 0.80  [0.45 – 1.42] | 0.82  [0.33 – 1.43] | 0.53  [0.09 – 1.20] | 0.81  [0.19 – 3.41] | 1.54  [1.16 – 1.73] | 1.90  [0.48 – 6.20] |
| **sEAS** | 0.38  [0.27 – 1.50] | 0.30  [0.20 – 1.36] | 0.92  [0.65 – 1.97] | 6.07  [0.52 – 9.23] | 1.08  [0.97 – 1.57] | 0.21  [0.12 – 2.33] |
| **maEAS** | 2.32  [1.31 – 3.85] | 1.82  [1.21 – 2.14] | 0.18  [0.08 – 0.54] | 0.74  [0.27 – 0.82] | 1.42  [1.27 – 1.5] | 1.92  [1.73 – 5.65] |

*sEAS= Severe Asthma, mEAS= Mild/Moderate Asthma, maEAS= Mastocytic Asthma, Ctr= Healthy

**Supplementary Table S5.** Cytokine concentrations of IL-5, IL-17A, IL-2, IL-4, IFNɣ, CXCL8, and TNFα measured via multiplex bead immunoassay in healthy (n=11), mild/moderate asthma (n = 9), and severe asthma (n = 14). Numbers in italics indicate this value was the software assigned lowest detectable concentration for the individual analyte. Data is represented at the median [IQR] for each group. sEAS= Severe Asthma, mEAS= Mild/Moderate Asthma, Ctr= Healthy.

| **Horse** | **IL-5**  **pg/mL** | **IL-17A**  **pg/mL** | **IL-2**  **pg/mL** | **IL-4**  **pg/mL** | **IFNɣ**  **pg/mL** | **CXCL8**  **pg/mL** | **TNFα**  **pg/mL** |
| --- | --- | --- | --- | --- | --- | --- | --- |
| **sEAS** |  |  |  |  |  |  |  |
| 1 | *0.5* | *0.49* | *0.44* | *16.35* | 81.24 | 46.66 | 91.34 |
| 2 | 0.92 | *0.49* | 0.46 | *16.35* | 31.69 | 16.86 | 4.92 |
| 3 | 28.38 | *0.49* | 20.66 | 120.22 | 981.27 | 84.69 | 111.55 |
| 4 | *0.5* | *0.49* | *0.44* | *16.35* | *21.08* | 20.99 | 1.43 |
| 5 | *0.5* | *0.49* | 0.55 | *16.35* | *21.08* | 7.49 | 1.43 |
| 6 | 2.2 | *0.49* | 0.46 | *16.35* | 26.11 | 19.68 | 4.41 |
| 7 | *0.5* | *0.49* | 0.46 | *16.35* | 154.19 | 38.91 | 20.45 |
| 8 | 1.6 | *0.49* | 0.46 | *16.35* | 56.72 | 20.99 | 25.72 |
| 9 | 1.46 | *0.49* | 2.48 | *16.35* | 73.52 | 19.97 | 17.93 |
| 10 | 1.05 | *0.49* | *0.44* | *16.35* | 28.87 | 24.12 | 15.04 |
| 11 | *0.5* | *0.49* | *0.44* | *16.35* | 38.87 | 16.86 | 5.3 |
| 12 | 3.02 | *0.49* | 0.86 | 19.98 | 49.2 | 19.54 | 14.03 |
| 13 | *0.5* | *0.49* | *0.44* | *16.35* | *21.08* | 9.9 | *0.8* |
| 14 | 1.9 | *0.49* | *0.44* | *16.35* | *21.08* | 18.65 | 8.59 |
| **Median [IQR]** | 0.99  [0.5 – 1.98] |  | 0.46  [0.44 – 0.63] | 16.35  [16.35 – 16.35] | 35.28  [21.08 – 75.45] | 19.83  [16.86 – 27.82] | 11.31  [3.67 – 21.77] |
| **mEAS** |  |  |  |  |  |  |  |
| 15 | 0.8 | *0.49* | *0.44* | *16.35* | 28.87 | 24.54 | 7.15 |
| 16 | 0.63 | *0.49* | *0.44* | *16.35* | *21.08* | 14.41 | 5.42 |
| 17 | 6.44 | *0.49* | 0.91 | 31.96 | 49.2 | 11.87 | 4.41 |
| 18 | 1.32 | *0.49* | *0.44* | *16.35* | 34.53 | 20.41 | 8.35 |
| 19 | 11.24 | *0.49* | 2.54 | 105.48 | 184.96 | 44.81 | 7.03 |
| 20 | 1.46 | *0.49* | *0.44* | *16.35* | *21.08* | 6.05 | 6.54 |
| 21 | *0.5* | *0.49* | *0.44* | *16.35* | 106.07 | 6.42 | 8.23 |
| 22 | 1.05 | *0.49* | 1.72 | *16.35* | *21.08* | 11.06 | 5.17 |
| 23 | 1.05 | *0.49* | *0.44* | *16.35* | *21.08* | 17.46 | 3.63 |
| **Median [IQR]** | 1.05  [0.72 – 3.95] |  | 0.44  [0.44 – 1.32] | 16.35  [16.35 – 24.16] | 28.87  [21.08 – 77.64] | 14.41  [8.74 – 22.48] | 6.54  [4.79 – 7.69] |
| **Ctr** |  |  |  |  |  |  |  |
| 24 | *0.5* | *0.49* | *0.44* | *16.35* | *21.08* | 1.6 | *0.8* |
| 25 | *0.5* | *0.49* | *0.44* | *16.35* | *21.08* | 1.6 | *0.8* |
| 26 | 0.99 | *0.49* | *0.44* | *16.35* | *21.08* | *1.26* | *0.8* |
| 27 | 3.02 | *0.49* | *0.44* | *16.35* | 28.87 | 12.03 | 2.28 |
| 28 | *0.5* | *0.49* | *0.44* | *16.35* | *21.08* | 6.42 | 4.41 |
| 29 | 0.86 | *0.49* | *0.44* | *16.35* | *21.08* | 9.56 | 2 |
| 30 | *0.5* | *0.49* | *0.44* | *16.35* | *21.08* | *1.26* | *0.8* |
| 31 | *0.5* | *0.49* | *0.44* | *16.35* | 22.04 | 6.78 | 4.15 |
| 32 | *0.5* | *0.49* | *0.44* | *16.35* | *21.08* | *1.26* | *0.8* |
| 33 | *0.5* | *0.49* | *0.44* | *16.35* | *21.08* | 8.88 | 3.63 |
| 34 | 3.02 | *0.49* | *0.44* | 28.99 | *21.08* | 4.54 | 3.1 |
| **Median [IQR]** | 0.5  [0.5 – 0.99] |  | 0.44  [0.44 – 0.44] | 16.35  [16.35 – 16.35] | 21.08  [21.08 – 21.08] | 4.54  [1.26 – 8.88] | 2.0  [0.8 – 3.63] |

**Supplementary Table S6.** Percent recovery of spiked samples.

| **Analyte** | **Expected (pg/mL)** | **Observed (pg/mL)** | **% Recovered** |
| --- | --- | --- | --- |
| **IL-5** | 3.66 | 4.49 | 122.68 |
|  | 14.65 | 15.06 | 102.80 |
|  | 58.59 | 62.36 | 106.43 |
|  | 234.38 | 238.25 | 101.65 |
|  | 937.5 | 866.24 | 92.40 |
|  | 3750 | 3336.18 | 88.96 |
| **IL-17A** | 3.66 | 2.37 | 64.75 |
|  | 14.65 | 5.65 | 59.04 |
|  | 58.59 | 37.58 | 64.14 |
|  | 234.38 | 143.81 | 61.36 |
|  | 937.5 | 543.24 | 57.95 |
|  | 3750 | 2023 | 53.95 |
| **IL-2** | 3.66 | 3.84 | 104.92 |
|  | 14.65 | 16.67 | 113.79 |
|  | 58.59 | 67.74 | 115.62 |
|  | 234.38 | 254.92 | 108.76 |
|  | 937.5 | 1058.77 | 112.94 |
|  | 3750 | 3955.77 | 105.49 |
| **IL-4** | 73.24 | 130.78 | 178.56 |
|  | 292.97 | 435.57 | 148.67 |
|  | 1172 | 1469 | 125.34 |
|  | 4688 | 5494 | 117.19 |
|  | 18750 | 17736 | 94.59 |
|  | 75000 | 71106 | 94.81 |
| **IFNɣ** | 122.07 | 133.4 | 109.28 |
|  | 488.28 | 513.67 | 105.2 |
|  | 1953 | 1942.76 | 99.48 |
|  | 7812 | 7485.76 | 95.82 |
|  | 31250 | 27647.76 | 88.47 |
|  | 125000 | 103586.76 | 82.87 |
| **CXCL8/IL-8** | 14.65 | 18.78 | 128.19 |
|  | 58.89 | 54.56 | 93.12 |
|  | 234.38 | 201.51 | 85.98 |
|  | 937.5 | 844.48 | 90.08 |
|  | 3750 | 3449 | 91.97 |
|  | 15000 | 8979 | 59.86 |
| **TNFα** | 0.98 | 2.82 | 287.75 |
|  | 3.91 | 4.99 | 127.62 |
|  | 15.62 | 18.54 | 118.69 |
|  | 62.5 | 66.74 | 106.78 |
|  | 250 | 270.67 | 108.27 |
|  | 1000 | 1056 | 105.6 |

**Supplementary Table S7.** Taqman Gene Expression Assay ID

| **Gene Target** | **Assay ID** |
| --- | --- |
| β-actin | Ec04176172_gH |
| CD117 | Ec03469720_m1 |
| Tryptase | Ec03468172_m1 |
| Chymase | ARMFXWU (custom) |
| Carboxypeptidase A3 | APU629N, ARH6ATU (custom) |
| Transforming growth factor-β1 | Ec03468030_m1 |
| Bone morphogenetic protein-7 | Ec04320876_m1 |

**Supplementary Figure S1.** Dilutional linearity as depicted by expected versus measured with linear regression analysis for all cytokines of interest: (a) IL-5, (b) IL-17A, (c) IL-2, (d) IL-4, (e) IFNɣ, (f) CXCL8, (g) TNFα. All show high linear dilution of spiked samples.

**(a)**

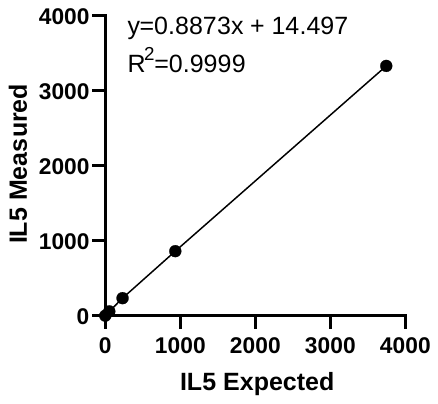

**(b)**

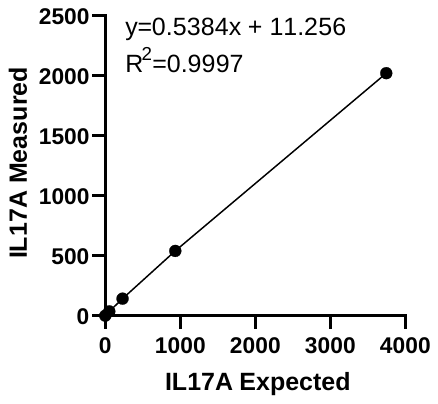

**(c)**

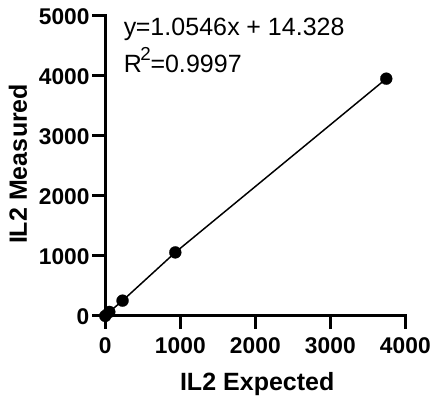

**(d)**

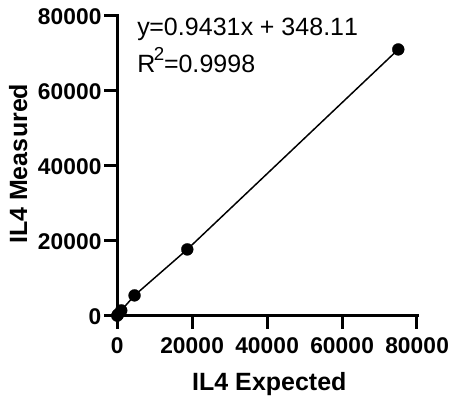

**(e)**

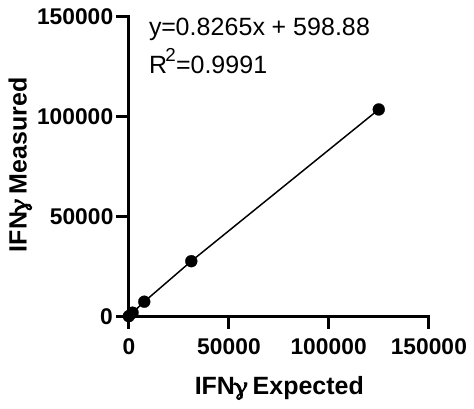

**(f)**

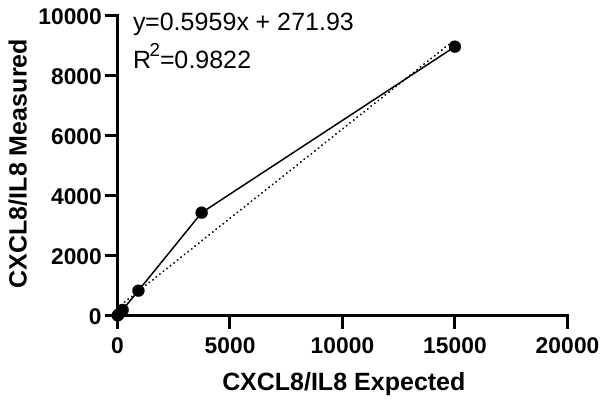

**(g)**

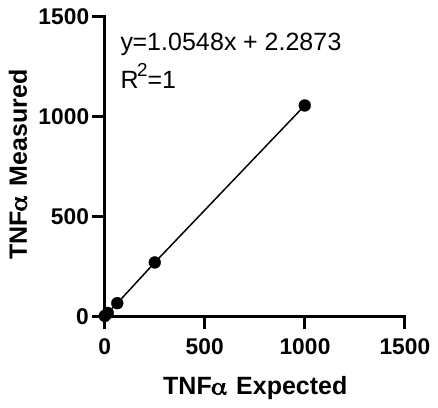
